# Roles of the 5-HT1A receptor in zebrafish responses to potential threat and in sociality

**DOI:** 10.1101/2024.04.17.588464

**Authors:** Loanne Valéria Xavier Bruce de Souza, Larissa Nunes Oliveira, Bruna Patrícia Dutra Costa, Monica Lima-Maximino, Vivianni Veloso, Caio Maximino

**Affiliations:** Laboratório de Neurociências e Comportamento “Frederico Guilherme Graeff”, Universidade Federal do Sul e Sudeste do Pará, Marabá/PA, Brazil; Programa de Pós-Graduação em Neurociência e Comportamento, Universidade Federal Pará, Belém/PA, Brazil; Rede de Biodiversidade e Biotecnologia da Amazônia Legal, Marabá/PA, Brazil; Laboratório de Neurofarmacologia e Biofísica – LaNeF, Universidade do Estado do Pará, Marabá/PA, Brazil

**Keywords:** zebrafish, model organism, heteroreceptor, vertebrates, defensive behavior

## Abstract

Anxiety is a normal emotion representing a reaction to potential danger, whereas fear can be defined as a reaction to real, explicit danger. Anxiety-like behavior in animal models has been associated with differences in the serotonergic system. Treatment of zebrafish cohorts with 8-OH-DPAT, a full agonist at the 5-HT1A receptor, decreased anxiety-like behavior in the novel tank test, but increased it in the phototaxis (light-dark preference) assay, both considered assays for anxiety-like behavior for this species. The same treatment decreased social approach in both the social investigation and social novelty phases of the social preference test. Blocking the 5-HT1A receptor with WAY 100,635 shifted the dose-response curve for the novel tank test rightward. These effects suggest a participation of the 5-HT 1A heteroreceptors in zebrafish anxiety and social preference, increasing anxiety and decreasing sociality. Thus, the study of this receptor is important for a better understanding of anxiety-like behavior in zebrafish and its relationship with similar phenomena in vertebrates.

Preprint: https://www.biorxiv.org/content/10.1101/2024.04.17.588464v1

Data: https://github.com/lanec-unifesspa/5HT/tree/main/5HT1A

**Lay summary:** This study looked at how a certain type of protein, the 5-HT1A receptor, affected anxiety in zebrafish. It’s important to understand that anxiety is a normal feeling we all get when we’re worried or scared about something, while fear is a reaction to an immediate danger. The researchers wanted to see how a specific drug that activates the 5-HT1A receptor, 8-OH-DPAT, affected the behavior of the fish. They used different tests to see how the fish reacted in different situations.

When the fish were given the drug, their anxiety-like behavior decreased in one test where they were put in a new tank. But in another test where they had to choose between light and dark areas, their anxiety-like behavior actually increased. It also made the fish less interested in being social with other fish. When a drug that blocks the 5-HT1A receptor, WAY 100,635, was give together with 8-OH-DPAT, evidence was found for a receptor reserve.

The researchers found that this drug affected the way the fish behaved in these situations. It seemed to make them more anxious in some cases and less interested in being social. This tells us that the receptor has an impact on how the fish are feeling, which can help us learn more about anxiety in zebrafish and, potentially, in other animals too, including humans.

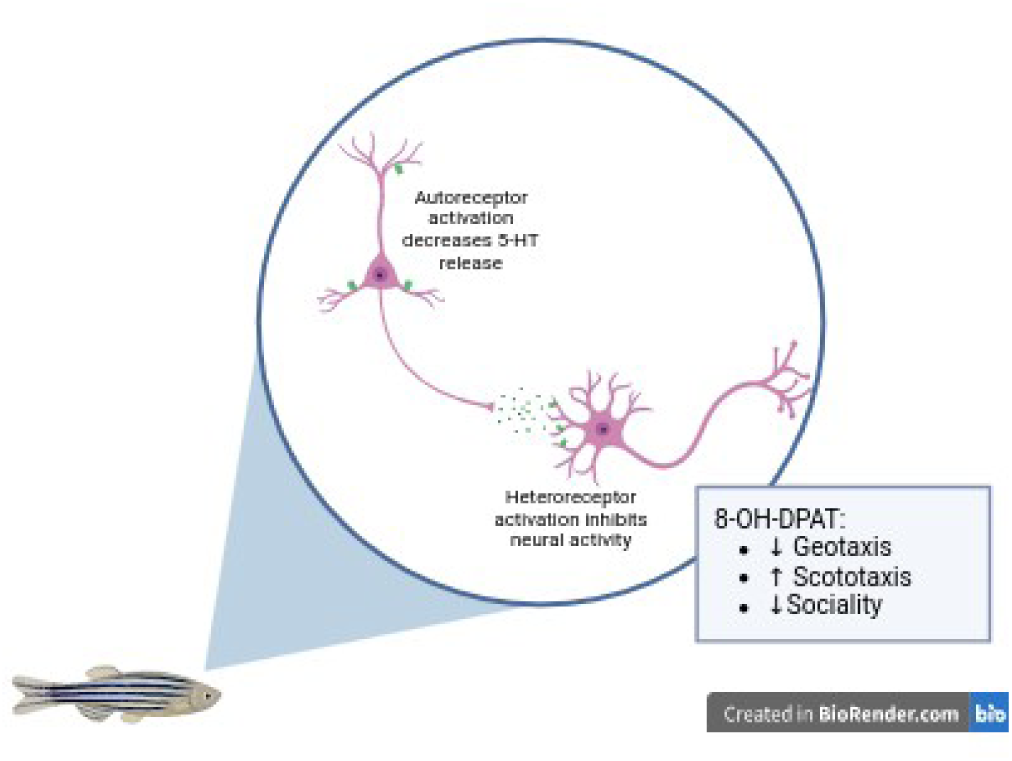

**Significance statement:** This research indicates that 5-HT1A heteroreceptors play a key role in increasing anxiety and decreasing sociality across vertebrates, shedding light on neural mechanisms that could inform treatments for anxiety-related disorders.

## 1. Introduction

Under situations of potential threat, humans and non-human animals adjust their behavior to cautious exploration, risk assessment, and behavioral inhibition, sometimes described as “anxiety-like behavior” (McNaughton & Corr, 2004). This pattern of defensive behavior is conceptually related to a fear-like state, but fear usually involves a response to an actual threat, whether that threat is proximal or distal (Perusini & Fanselow, 2015). While both are part of a taxonomy of defensive behavior, there is growing evidence to show that anxiety and fear are distinct emotions in terms of behavioral responses, physiological settings, and neural bases (Fanselow & Lester, 1988; McNaughton & Corr, 2022; McNaughton & Zangrossi Jr, 2008; Perusini & Fanselow, 2015).

The relationship between the neurobehavioral responses to threat and the serotonergic system points to a complex regulation, in which different subpopulations of serotonergic neurons in the dorsal raphe nucleus and the medial raphe nucleus are uniquely involved in the pathophysiology of anxiety and affective disorders (Deakin & Graeff, 1991; Maximino, 2012; Paul et al., 2014). This control is thought to be mediated by topographically organized projections to different areas of the limbic forebrain and midbrain (Abrams et al., 2004, 2005; Paul & Lowry, 2013). Serotonergic projections to forebrain regions, such as the amygaloid nuclei and the ventral hippocampus, increase behavioral responses to potential threat when activated, possibly via 5-HT2 and 5-HT3 receptors (Graeff et al., 1997; Maximino, 2012; Olivier et al., 2000; Quesseveur et al., 2012). In the midbrain – especially in the periaqueductal gray area –, activation of 5-HT1A and 5-HT2 receptors lead to an inhibition of neurobehavioral responses to actual threat (Castilho et al., 2002; Gomes & Nunes-de-Souza, 2009; Graeff, 2004; Graeff et al., 1997; Lima-Maximino et al., 2020; Paul et al., 2014). Thus, the Deakin-Graeff theory postulates not only the apparently paradoxical “double role” of serotonin on fear-like and anxiety-like behaviors in relation to different parts of the neuraxis, but also the differential participation of serotonin receptors in these effects.

The 5-HT1A receptor (5-HT1AR) is one of the most abundant and widely expressed receptors in the brain and the main inhibitory serotonergic receptor (Akimova et al., 2009; Albert & Vahid-Ansari, 2019). This receptor is found in two populations, the presynaptic autoreceptors (which promote local inhibitory control of neuronal firing rates and 5-HT synthesis in serotonergic neurons), which are located in the cell body and dendrites of 5-HT neurons; and postsynaptic heteroreceptors (which modulate the release of other neurotransmitters in non-serotoninergic neurons), which are located in the dendrites and cell body of non-serotoninergic neurons in 5-HT projection areas (Albert & Vahid-Ansari, 2019).

There is some evidence for a large receptor reserve for autoreceptors in opposition to heteroreceptors (Cox et al., 1993; Meller et al., 1990), which could explain why azapirones, partial agonists of the 5-HT1AR with anxiolytic-like activity in clinical settings (Chessick et al., 2006), act as antagonists at heteroreceptor sites and full agonists in autoreceptors (Newman-Tancredi et al., 2019). Low doses of 8-OH-DPAT, a full agonist, and buspirone, a partial agonist, preferentially activate 5-HT1A autoreceptors, and consequently, the release of 5-HT from serotonergic nerve endings decreases. Therefore, 5-HT synthesis is reduced as a feedback mechanism. From a systemic point of view, therefore, the effect of buspirone leads to inhibition of the activity of serotonergic neurons (and consequent decrease in the release of 5-HT) associated with the blockade of 5-HT1A postsynaptic receptors. Full agonists, such as 8-OH-DPAT, appear to act as full agonists on both autoreceptors and heteroreceptors (Sprouse & Aghajanian, 1988), decreasing the availability of 5-HT in other receptor subtypes at the same time that post-synaptic 5-HT1ARs are activated. This discrepancy could explain why both partial agonists and antagonists of the 5-HT1AR decrease defensive behavior to potential threat (Cao & Rodgers, 1997; Maximino et al., 2013; Rodgers & Cao, 1997). The putative differential effects of drugs on auto- and heteroreceptors lead to the necessity of understanding the roles of these different sites in the control of defensive behavior. Studies using mice lacking 5-HT1A auto- or heteroreceptors (Richardson-Jones et al., 2011), 5-HT1AR knockouts with expression rescued in pre- or post-synaptic sites (Gross et al., 2002; Piszczek et al., 2013), intact animals treated with biased agonists that activate pre- or postsynaptic 5-HT1ARs *in vivo* (Jastrzębska-Więsek et al., 2018), or microinjection of agonists in the raphe or in postsynaptic sites (File et al., 1996; File & Gonzalez, 1996), produced conflicting results.

The roles of 5-HT1ARs were also investigated in non-mammalian model organisms, including zebrafish (*Danio rerio* Hamilton 1822). This model organism has increasingly been used in neuroscience and behavioral pharmacology to understand the mechanisms of defensive behavior (Cueto-Escobedo et al., 2022; Gerlai, 2020, 2023; MacRae & Peterson, 2023). While there is considerable conservation of the serotonergic system is in zebrafish, there are important differences in the degree of neuroanatomical and genomic conservation (Herculano & Maximino, 2014). For example, zebrafish present extra serotonergic nuclei in the brain in addition to the raphe (Kaslin & Panula, 2001; Lillesaar et al., 2009), and a genomic duplication event led to multiple copies of genes involved in serotonin synthesis, transport, and metabolism, as well as of serotonin receptors (including two copies of the 5-HT1AR) (Norton et al., 2008; Sourbron et al., 2016). The two copies of the 5-HT1AR-coding gene, *htr1aa* and *htr1ab*, are expressed in the adult zebrafish brain in a complementary manner, with high expression of the *htr1aa* isoform as an autoreceptor in the pretectal diencephalic cluster, superior raphe nucleus, and paraventricular organ, while the *htr1ab* isoform is expressed as an autoreceptor in the caudal zone of the periventricular hypothalamus, paraventricular organ, superior raphe, and pretectal cluster (Norton et al., 2008). As an heteroreceptor, *htr1aa* is found in the ventral nucleus of the ventral telencephalic area (Vv, homologous to the septal nuclei), preoptic region (PPa), periventricular gray zone of the optic tectum (PGZ), posterior tuberculum, central posterior thalamic nucleus, paraventricular organ, and ventral zone of the periventricular hypothalamus (Hv), while *htr1ab* is found in central gray (homologous to the periaqueductal gray area), Hv, zona limitans, PGZ, and PPa (Norton et al., 2008). Zebrafish exposed to the phototaxis assay (light-dark test), a commonly used test for anxiety-like behavior in this species, show increased *cfos* expression in regions with high density of 5-HT1AR isoforms, such as Vv and Hv (Lau et al., 2011).

Despite the differences in genes and expression of 5-HT1ARs in zebrafish and mammals, the role of the serotonergic system appears to be relatively conserved in zebrafish (Herculano & Maximino, 2014). Some evidence suggests a role for 5-HT1A receptors in anxiety-like behavior in zebrafish; for example, buspirone produces consistent type-anxiolytic effects in behavioral assays with zebrafish, regardless of the route of administration, dose, or test (Abozaid & Gerlai, 2022; Bencan et al., 2009; Connors et al., 2014; Facchin et al., 2015; Gebauer et al., 2011; Hawkey et al., 2021; Lau et al., 2011; Maaswinkel et al., 2012, 2013; Maximino et al., 2011, 2014). Maximino et al. (2013) evaluated behavior characterized by scototaxis (preference for dark environments over light environments) and geotaxis (the measurement of time at the top or bottom of the apparatus). They observed anxiety-like behavior correlated positively with extracellular levels of 5-HT in the brain in the novel tank test (NTT), while the correlation was negative in the phototaxis assay. This suggests that 5-HT may regulate zebrafish behavior differently in the NTT and phototaxis assay. Nonetheless, buspirone produced an anxiolytic-like effect in both the NTT and the phototaxis assay. Buspirone also has been shown time spent near a conspecific in a social investigation test, but not in the social novelty test (Barba-Escobedo & Gould, 2012), which could either represent an anxiolytic-like effect or increased social motivation. Puzzlingly, WAY 100635, a full 5-HT1A receptor antagonist, also produced anxiolytic-like effects in both the NTT and the phototaxis assay (Maximino et al., 2013). Nowicki et al. (2014), on the other hand, observed an anxiogenic-like effect on the NTT after treatment with p-MPPF, another 5-HT_1A_ receptor antagonist. On the other hand, treatment of zebrafish with WAY 100,635 increased responsiveness to an olfactory-evoked fear stimulus (conspecific alarm substance) in this species (Nathan et al., 2015), suggesting opponent roles of this receptor in responses to potential vs. actual threat (anxiety vs. fear) in this species.

The similar effects of buspirone, a partial agonist, and WAY 100,635, a full antagonist, on zebrafish anxiety-like behavior suggest the existence of a receptor reserve in presynaptic sites. Furthermore, there are no studies with full agonists in this species. Thus, the aim of the present study is to evaluate the effect of treatment with 8-OH-DPAT, a full agonist of the 5-HT1A receptor, on behaviors relevant to anxiety and sociability in zebrafish. 8-OH-DPAT is a full agonist with high affinity for the 5-HT1A receptor (pKi for human receptors between 8.4 and 9.4; https://www.guidetopharmacology.org/GRAC/LigandDisplayForward?tab=biology&ligandId=7), although it has been reported to act as a full agonist at 5-HT7 receptors as well (pKi for rat receptors between 7.3 and 7.5; https://www.guidetopharmacology.org/GRAC/LigandDisplayForward?tab=biology&ligandId=7). We hypothesized that treatment with 8-OH-DPAT would be anxiogenic, and predicted that this treatment would reduce time on white and increase risk assessment in the phototaxis assay, increase time on bottom and erratic swimming in the NTT, and decrease preference scores in both the social investigation and social novelty tests. Moreover, given the evidence for a receptor reserve in mammals (Cox et al., 1993; Meller et al., 1990), we hypothesized that pre-treatment with the antagonist WAY 100,635 would displace the dose-response curve rightward.

## 2. Materials and methods

### 2.1 Animals and housing

96 adult zebrafish (*Danio rerio*) (>4 months old) from the shortfin phenotype were used in the experiments. Adult wild-caught zebrafish (4-6 months old) of both sexes from the strain were obtained from a colony resulting from cross-breeding between animals purchased in specialized stores (Power Fish Fisheries, Itaguaí/RJ, Brazil). Based on body morphology (general body shape and presence of genital papilla in females (Parichy et al., 2009), a male:female ratio of approximately 40%:60% was estimated; sex of samples was ultimately established after animals were sacrificed, using histological analysis of the gonads (Siegfried & Steinfeld, 2021). All animals were kept in 50L tanks and acclimatized in the laboratory for a period of 14 days before the onset of experiments. The tanks were kept constant at a temperature of 26 ± 2°C and pH 7.2, oxygenation, 14/10 light-dark cycle provided by fluorescent lamps, according to standards of care for zebrafish (Lawrence, 2007). Animals were fed twice a day with commercial flocked feed. Animal density adopted was in accordance with the protocol adopted in the Brazilian guideline for the care of zebrafish in the laboratory (Conselho Nacional de Controle de Experimentação Animal - CONCEA, 2017). The project was carried out with the approval of the Animal Use Ethics Committee of UNIFESSPA (Universidade Federal do Sul e do Sudeste do Pará) protocol number 23479.019576/2022-43.

Sample sizes were calculated based on a model for independent-samples t-tests, considering an effect size of *d* = 1.35 (based on a metanalysis of the effects of buspirone on behavioral tests in zebrafish; https://github.com/lanec-unifesspa/5HT/blob/main/5HT1A/zf_metanalysis_buspirone.pdf), a power of 0.8, and α = 0.05. In these conditions, a minimum sample size of *n* = 10 animals/group was projected; ultimately, 12 animals per group were used in each experiment. Calculations were made using the R package pwr (v. 1.3-0; https://cran.r-project.org/web/packages/pwr/index.html).

### 2.2 Drugs and treatments

8-OH-DPAT (8-hydroxy-2-(dipropylamino)tetralin; CAS#87394-87-4) and WAY 100,635 (*N*-[2-[4-(2-Methoxyphenyl)-1-piperazinyl]ethyl]-N-(2-pyridyl)cyclohexanecarboxamide) were bought from Sigma-Aldrich (St Louis, USA) and dissolved in Cortland’s salt solution (NaCl 124.1 mM, KCl 5.1 mM, Na2HPO4 2.9 Mm, MgSO4 1.9 mM, CaCl2 1.4 mM, NaHCO3 11.9 mM, 1000 units heparin; Perry et al., 1984). For all treatments, the solutions were injected intraperitoneally into cold-anesthetized, and once anesthetized, they were positioned on a sponge-based surgical bed for injection according to the method described by Kinkel et al. (2010). For both treatments, the injection was carried out using a microsyringe (Hamilton 701N, 26 gauge needle, 200 conical tip, 10 μL), with a dose proportional to the animal’s body weight (0.3 mg/kg) and an injection volume of 5 μL.

Experimenters were blind to treatment through the use of coded vials; only the principal investigator had access to the codes before data analysis. Treatment order was randomized within sessions by using a random number generator (https://www.randomizer.org/).

### 2.3. Experiment 1: Novel tank test

In general, this task aims to evaluate zebrafish vertical locomotion/exploration as an index of defensive responses related to the response to novelty (Bencan et al., 2009; Egan et al., 2009; Levin et al., 2007). Behavioral assessment of the animals was carried out according to the methodology previously described in works by the research group (https://dx.doi.org/10.17504/protocols.io.bp2l6x45klqe/v1), in which parameters related to locomotion and the vertical position occupied by the animal in the tank (15 cm x 25 cm x 20 cm; width, length, height) were evaluated. Dechlorinated water was used for both the recovery tank and the test tank. 12 adult zebrafish were used per group; the same 24 animals were also used in Experiment 2. Immediately after the intraperitoneal injection, the fish were relocated with a fish net to a recovery tank, where they remained for 30 minutes. Then, using the fish net, the fish was transferred to the test tank, where it remained for the 6-min habituation period, during which time the fish’s behavior was recorded, generating a spatio-temporal exploratory profile pattern. The behavioral endpoints were transcribed using The Real Fish Tracker software (v. 0.4.0; https://www.dgp.toronto.edu/~mccrae/projects/FishTracker/). Table 1 summarizes the endpoints assessed in Experiment 1.

**Table 1.**
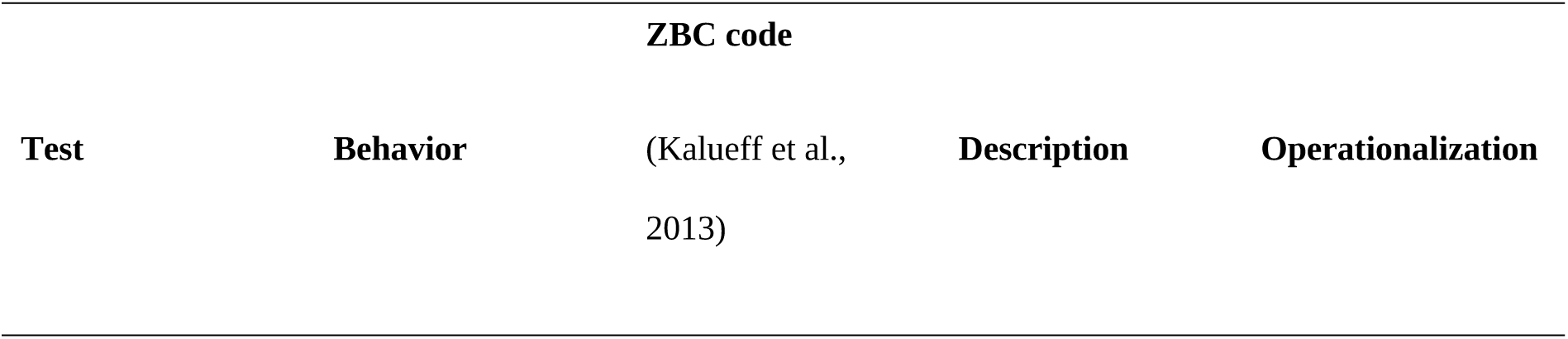
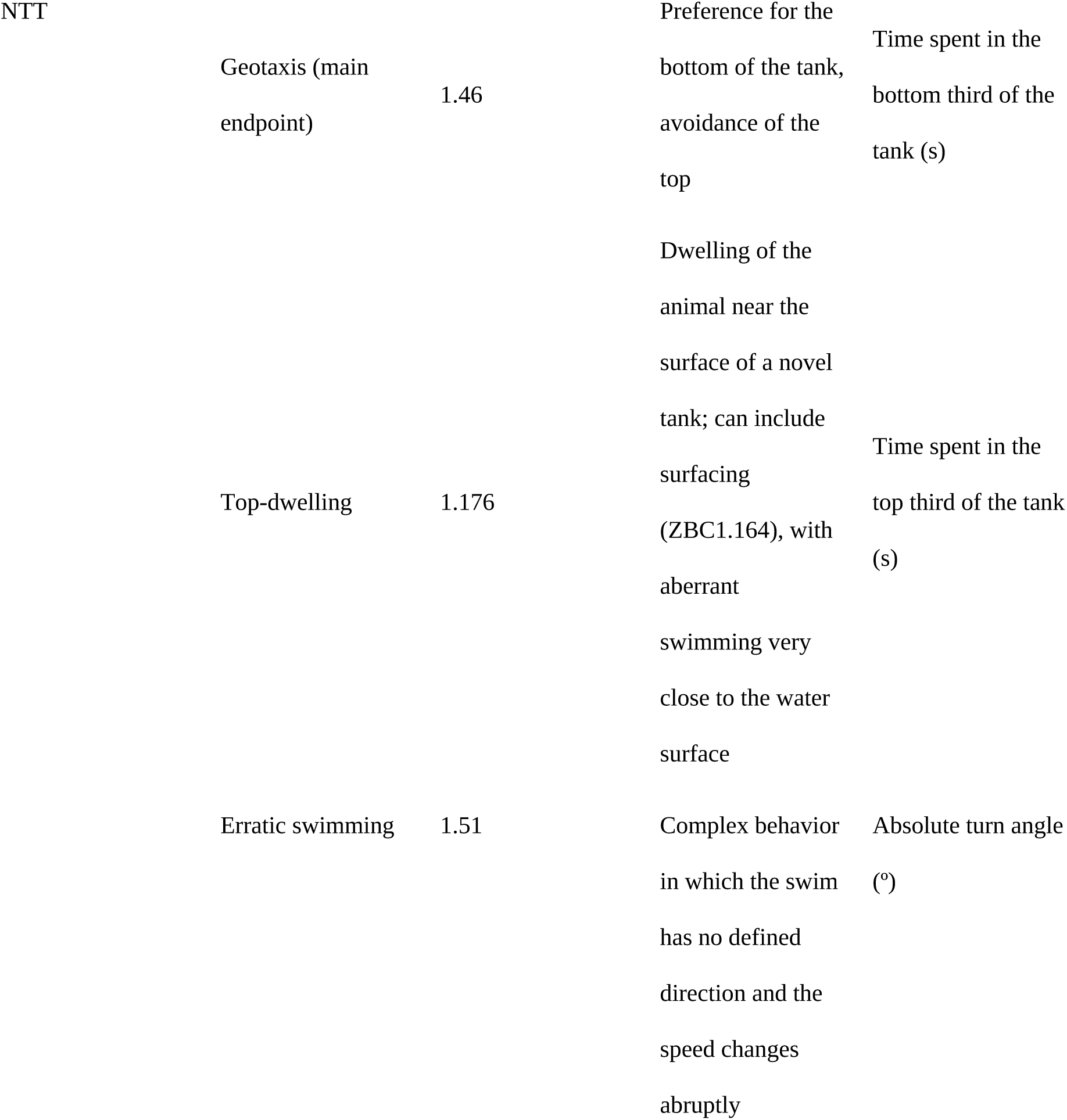
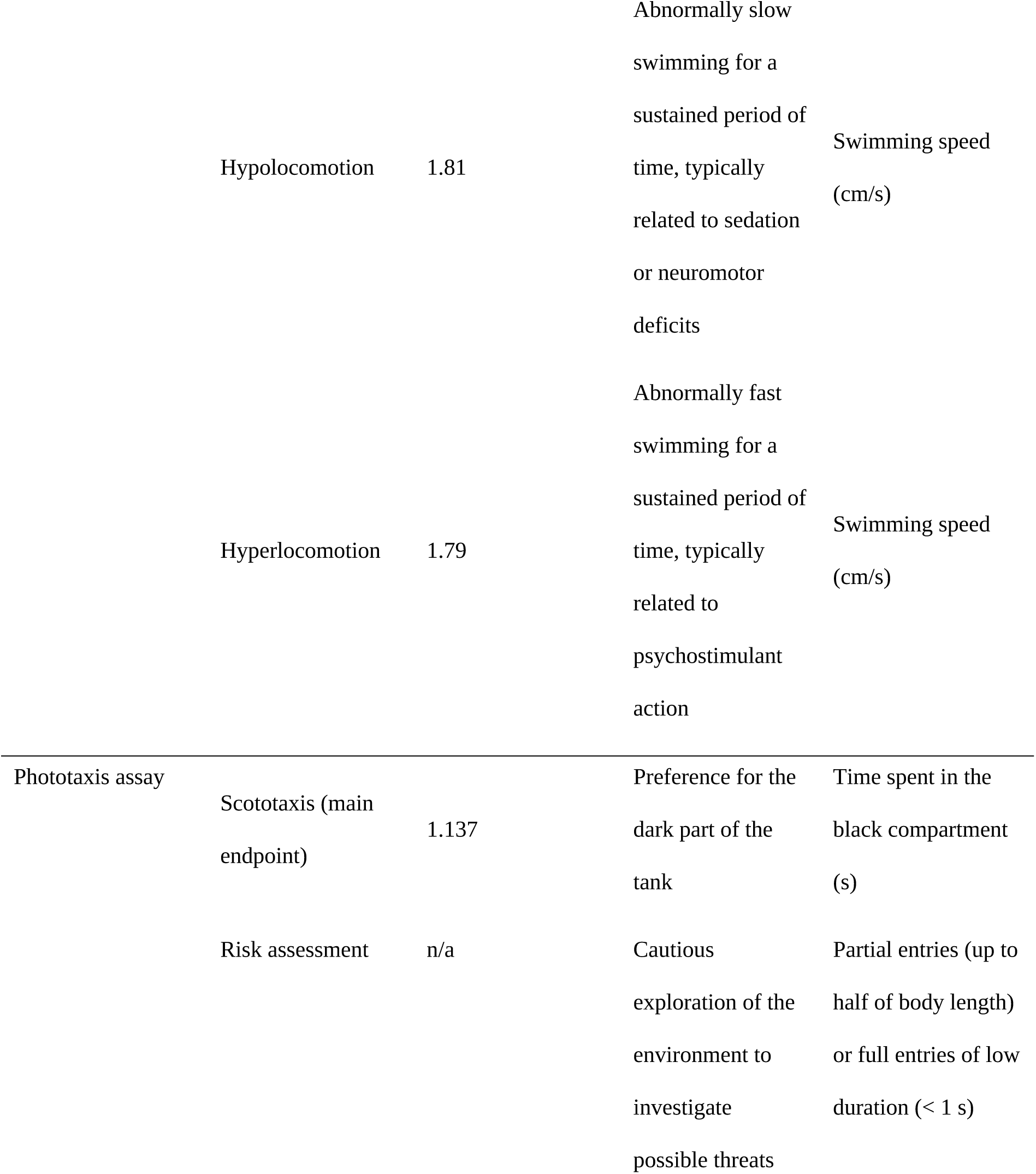
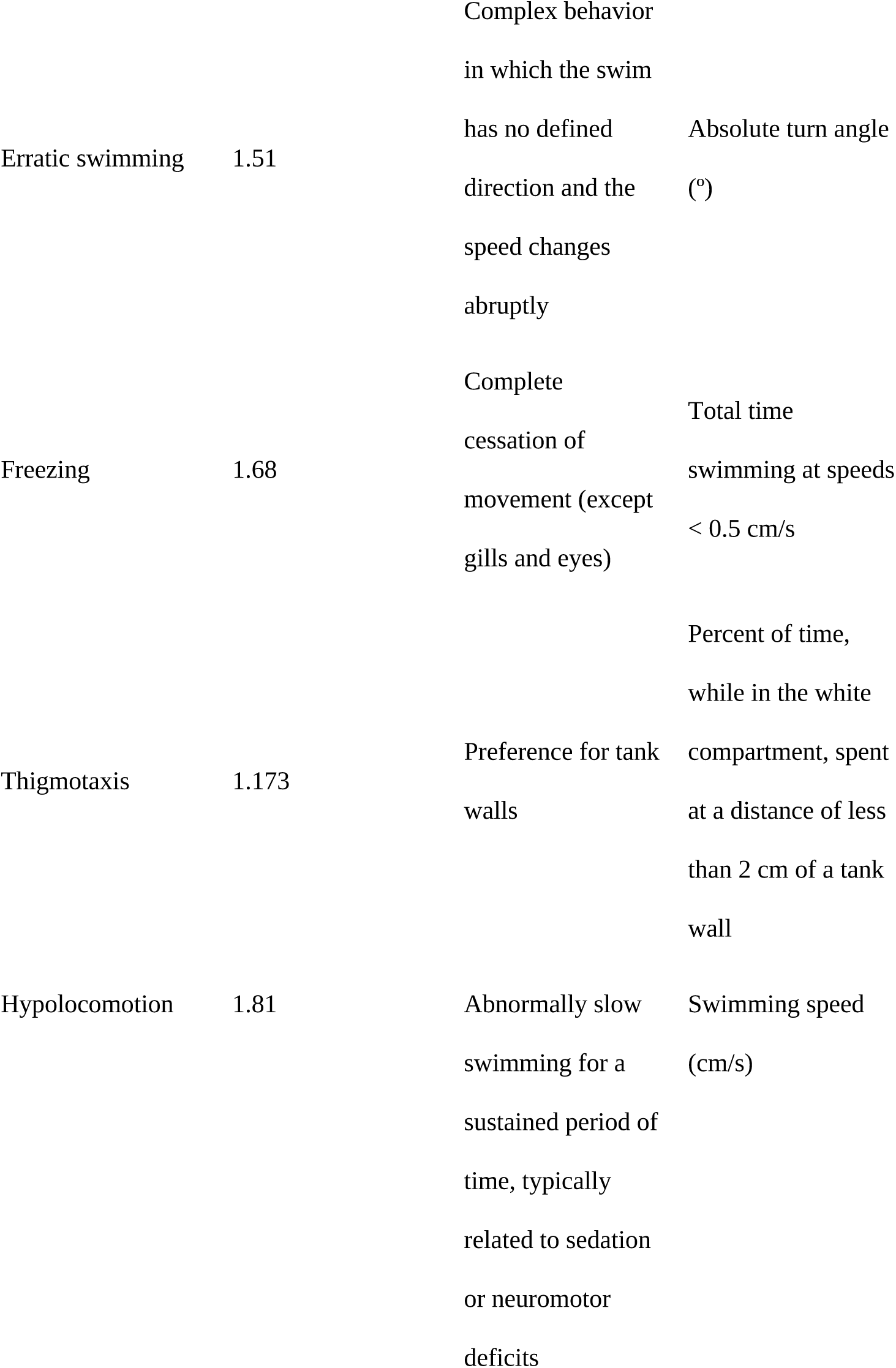
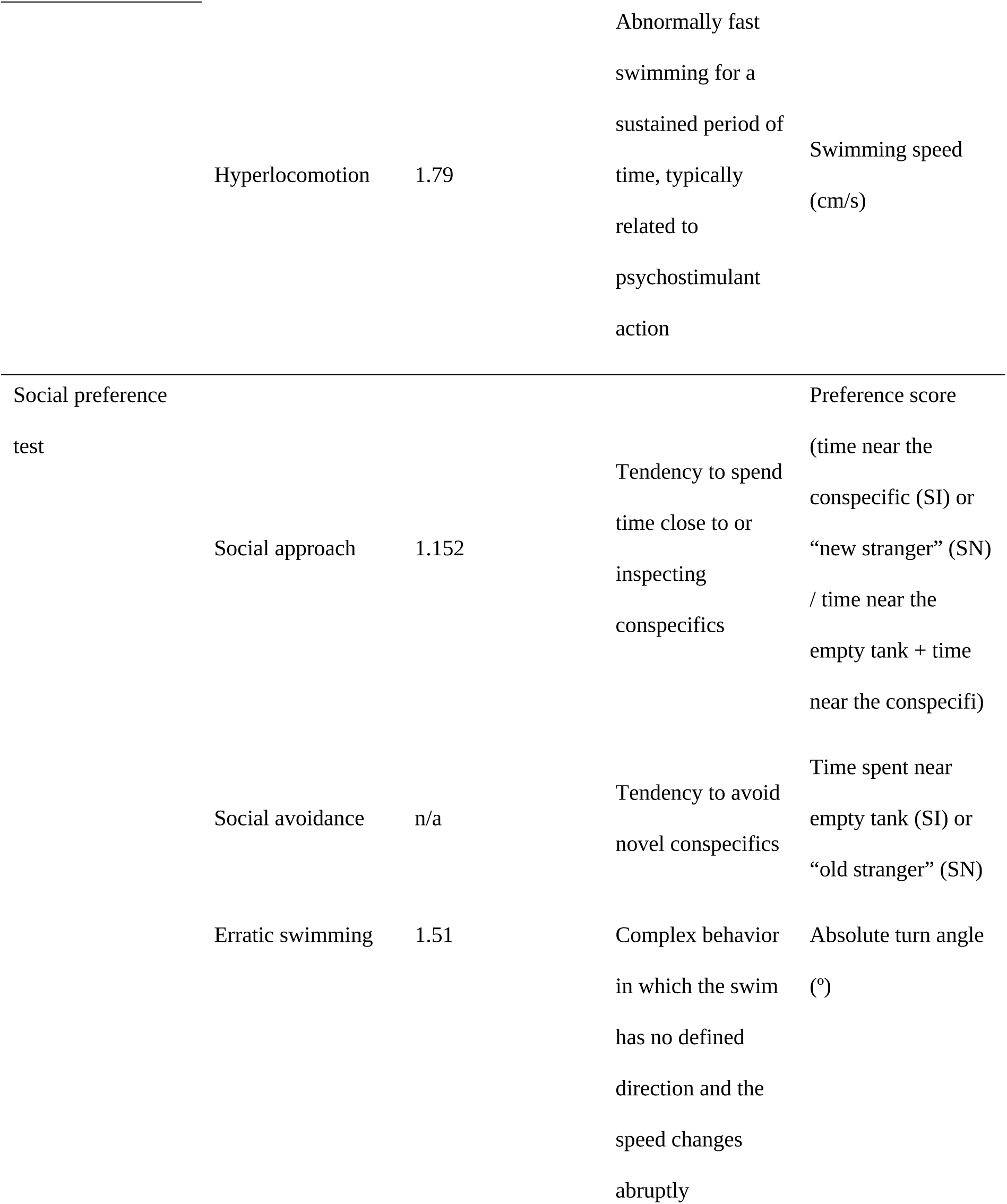
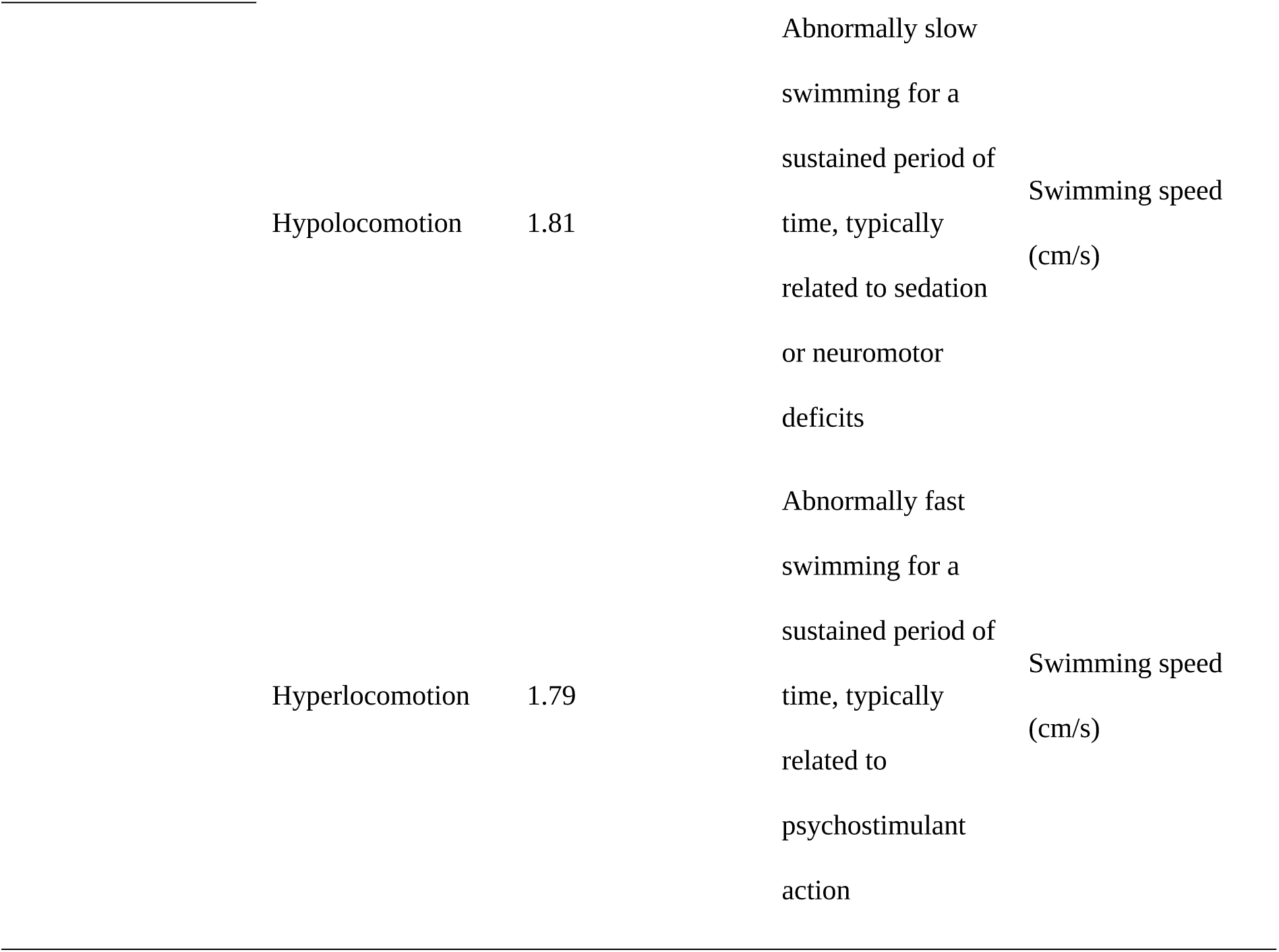
Behavioral endpoints assessed in the novel tank test (NTT), phototaxis assay, and social preference.

### 2.4. Experiment 2: Phototaxis assay

Light/dark preference in zebrafish, characterized by a preference for either dark or light environments, has been described and validated as a model for evaluating anxiety-like behavior for the species (Araujo et al., 2012; Maximino et al., 2010, 2011; Serra et al., 1999). Preference for either light or dark environments have been described in the literature, depending on developmental stage (e.g., more preference for light environments in larvae and juvenile zebrafish; Chen et al., 2015; Steenbergen et al., 2011) and light levels (e.g., more preference for dark/black environments under bright light; Araujo et al., 2012; Blaser & Peñalosa, 2011; Facciol et al., 2017, 2019). In both cases, anxiolytic drugs, including buspirone, have been shown to reduce preference (Maximino et al., 2011; Steenbergen et al., 2011). 12 zebrafish were used in each group. Since animals were reused in Experiments 1 and 2, the order in which animals were subjected to either the NTT or the phototaxis assay was randomized. Immediately after the intraperitoneal injection, the fish was relocated with an fish net to a recovery tank, where it remained for 30 min. Then, using the fish net, the fish was transferred to an apparatus (15 cm x 10 cm x 45 cm; height, width, length) divided into a white compartment and a black compartment, with hatches in the middle where the fish is placed for acclimatization for 3 min. After this, the hatches were removed and the fish was free to explore the apparatus for 15 min. The protocol established in the research group (https://dx.doi.org/10.17504/protocols.io.puydnxw) was used. Scototaxis, swimming speed, erratic swimming, and freezing were extracted using TheRealFishTracker (v. 0.4.0; https://www.dgp.toronto.edu/~mccrae/projects/FishTracker/), while risk assessment and thigmotaxis were scored manually with the help of X-Plo-Rat (Tejada et al., 2018; https://sites.ffclrp.usp.br/scotty/). Table 1 summarizes the endpoints assessed in Experiment 2.

### 2.5. Experimental 3: Social preference test

The social preference test was divided in two stages, social interaction and social novelty tests, and carried out based on the protocol described by Barba-Escobedo and Gould (2012). These tasks aim to analyze the tendency of zebrafish to approach co-specifics and investigate novel social situations. 48 adult zebrafish were used as conspecific stimuli, while 24 adult zebrafish were used as focal fish in the test, where 12 animals were injected intraperitoneally with the vehicle solution (Cortland’s saline solution), and 12 animals with 8-OH-DPAT. Animals from this experiment were not reused from Experiments 1 and 2, due to the longer extension of the test; exposing animals to three behavioral tests would risk the drug having been eliminated by the end of the third task. Conspecifics were drawn from different tanks than the hometank of the focal fish, and therefore focal fish had no prior contact with them. Immediately after intraperitoneal injection, the focal fish was relocated using a fish net to a recovery tank, where it remained for 30 min. Then, using the fish net, the focal fish is transferred to the test tank where it remains for 3 min for acclimatization. For both the social interaction and social novelty tests, three tanks were used: the main tank (15 cm x 25 cm x 20 cm; width, length, height), for the focal animal, and two other tanks for the conspecifics (15 cm x 17 cm x 16 cm; width, length, height). Both sides of the main tank were separated from the lateral tanks with barriers to prevent visual access. Once the fish’s acclimatization period was over, the barriers were removed allowing the focal fish visual access to the conspecifics. In the social interaction test (SI), the behavior of 1 focal animal was evaluated for an initial period of 6 min, in which it will have visual access to an unknown conspecific and an empty tank. After that time interval, barriers were lowered, a new conspecific was transferred to the empty tank, and the barriers were again removed to allow the focal fish visual access to the conspecifics. The behavior was observed again for 6 minutes, consisting in the social novelty test (SN). The conspecific that was present in the former stage was labeled as “old stranger”, while the conspecific that was introduced in the empty tank was labeled as “new stranger”. In both stages, the conspecific stimulus was a female zebrafish, confirmed by body morphology and, later, gonadal histology; female zebrafish stimuli are reported to produce stronger preferences than male stimuli (Ogi et al., 2021). In both the social interaction and social novelty stages, behavior was filmed and later analyzed using TheRealFishTracker software (v. 0.4.0; https://www.dgp.toronto.edu/~mccrae/projects/FishTracker/); behavioral endpoints are described in Table 1.

### 2.6. Experiment 4: Evidence for a receptor reserve

For Experiment 4, animals were first pre-treated with either vehicle or WAY 100,635 (0.001 mg/kg, a dose below the necessary to produce behavioral effects in adult zebrafish; Maximino et al., 2013; Nathan et al., 2015), and then injected with 8-OH-DPAT at the following doses: 0.03 mg/kg, 0.3 mg/kg, and 3 mg/kg. 30 min. after injection, animals were subjected to the novel tank test, as described in Section 2.3, above.

### 2.7. Statistical analysis

Results obtained from the behavioral tests from Experiment 1-3 were analyzed using unpaired t-tests or Wilcoxon rank sum test with continuity correction for variables which violated the assumptions of parametric statistics. No data was removed from the dataset. For Experiment 4, data were analyzed using two-way (8-OH-DPAT dose X WAY 100,635 dose) analyses of variance. The data was represented as Gardner-Altman plots (Ho et al., 2019), with individual datapoints plotted on the upper axes. On the lower axes, each effect size (Cohen’s d; Cohen, 1988) was plotted as a bootstrap sampling distribution. Bootstrap resamples were taken using Efron’s (1987) method; the confidence interval is bias-corrected and accelerated. Plots were made using R 4.1.2, using the dabestr package (v. 2023.9.12; https://github.com/ACCLAB/dabestr). Post-hoc power calculations were made using the pwr package (v. 1.3-0; https://cran.r-project.org/web/packages/pwr/index.html) in R, and the power of each analysis is reported. While sex was not included as a variable in the analysis, as the study was not designed to understand sex differences in the effect of 8-OH-DPAT, exploratory analyses were made on the data, following Joel and Fausto-Sterling (2016). Potential differences between males and females were explored via interaction effect size plots. Plots for these exploratory analyses were made using the esci package (https://github.com/rcalinjageman/esci) in jamovi (v. 2.4; https://www.jamovi.org).

### 2.7. Open science practices

Hypotheses were not formally preregistered. Data packages and analysis scripts can be found on our GitHub repository (https://github.com/lanec-unifesspa/5HT/tree/main/5HT1A).

## 3. Results

### 3.1. Novel tank test

8-OH-DPAT decreased geotaxis (t_[df = 19.484]_ = 5.222, p < 0.001, power = 0.999; Figure 1A) and increased top-dwelling (t_[df = 16.386]_ = −4.009, p < 0.001, power = 0.97; Figure 1B). No effects were observed on erratic swimming (t_[df = 21.998]_ = −0.159, p = 0.875, power = 0.05; Figure 1C), freezing (t_[df = 17.214]_ = −1.555, p = 0.138, power = 0.31; Figure 1D), or swimming speed (t_[df = 20.107]_ = 0.971, p = 0.343, power = 0.16; Figure 1E). For all variables, 95% confidence intervals for the effect sizes for sex X group interactions overlapped zero (Supplemental File S1); however, plots suggest higher geotaxis and lower top-dwelling in males, and larger effects of 8-OH-DPAT in females vs. male zebrafish in geotaxis and top-dwelling.

**Figure 1.**
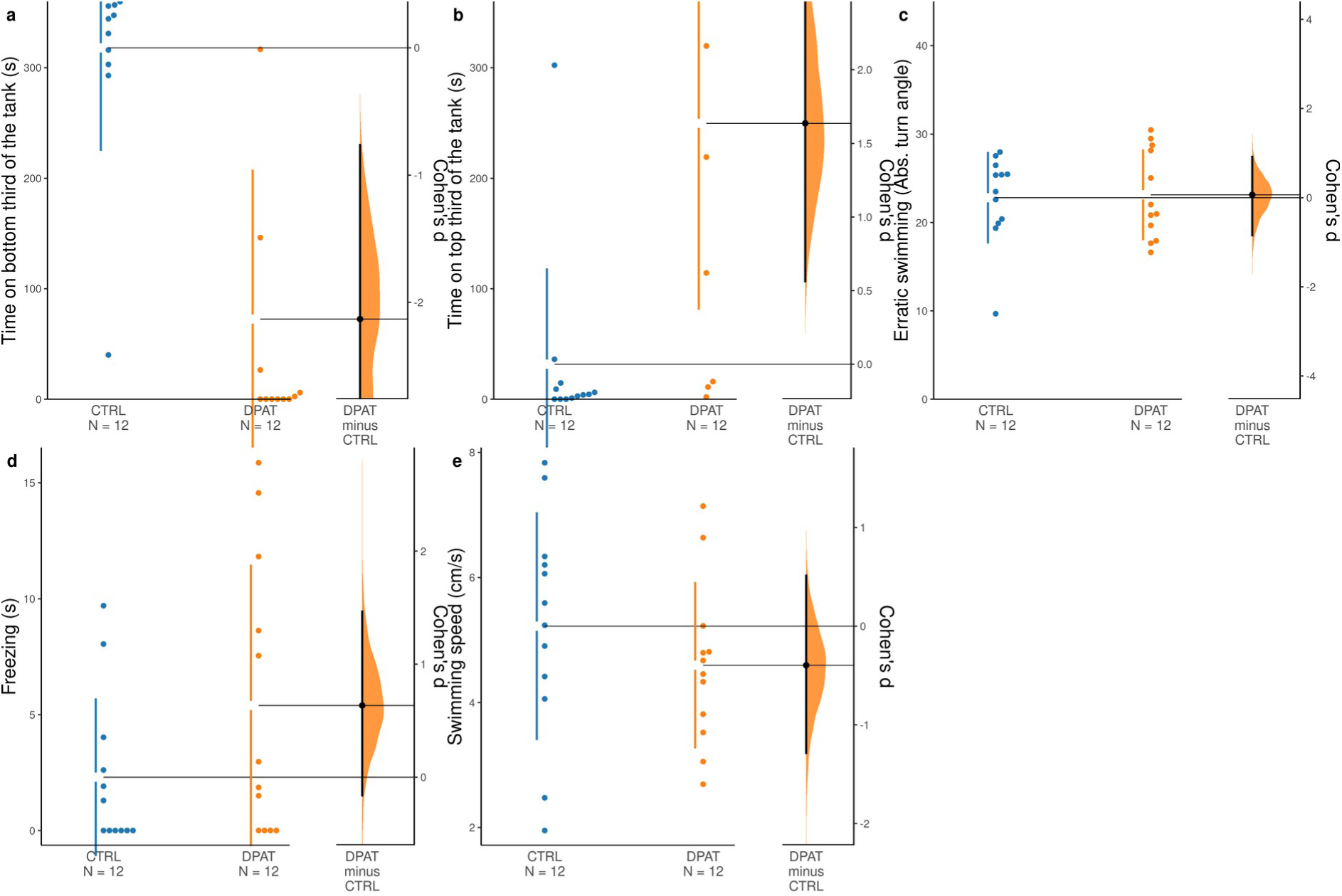
– 8-OH-DPAT decreases anxiety-like behavior in the zebrafish novel tank test. (A) Geotaxis. (B) Top-dwelling. (C) Erratic swimming. (D) Freezing. (E) Swimming speed. For a description of variables, refer to Table 1. Blue dots represent animals treated with vehicle (CTRL), and orange dots represent animals treated with 8-OH-DPAT (DPAT). The Cohen’s *d* between CTRL and DPAT is shown in Gardner-Altman estimation plot. Both groups are plotted on the left axes; the effect size (Cohen’s *d*) is plotted on floating axes on the right as a bootstrap sampling distribution. The mean difference is depicted as a dot; the 95% confidence interval is indicated by the ends of the vertical error bar. 5000 bootstrap samples were taken; the confidence interval is bias-corrected and accelerated.

### 3.2. Phototaxis assay

8-OH-DPAT increased scototaxis (t_[df = 17.361]_ = 2.43, p = 0.262, power = 0.64; Figure 2A) and increased risk assessment (W = 33, p = 0.024, power = 0.6; Figure 2B). No effect were found for erratic swimming (t_[df = 20.671]_ = 0.727, p = 0.475, power = 0.11; Figure 2C), freezing (W = 71.5, p = 0.99, power = 0.06; Figure 2D), thigmotaxis (t_[df = 14.873]_ = −1.323, p = 0.206, power = 0.24; Figure 2E), or swimming speed (t_[df = 21.985]_ = 1.71, p = 0.101, power = 0.37; Figure 2F). For all variables, 95% confidence intervals for the effect sizes for sex X group interactions overlapped zero (Supplemental File S2); however, plots suggest higher scototaxis in females, and larger effects of 8-OH-DPAT in males vs. female zebrafish in this variable; as well as higher thigmotaxis in males, with a small-sized interaction effect in which 8-OH-DPAT increases thigmotaxis in males but not females.

**Figure 2.**
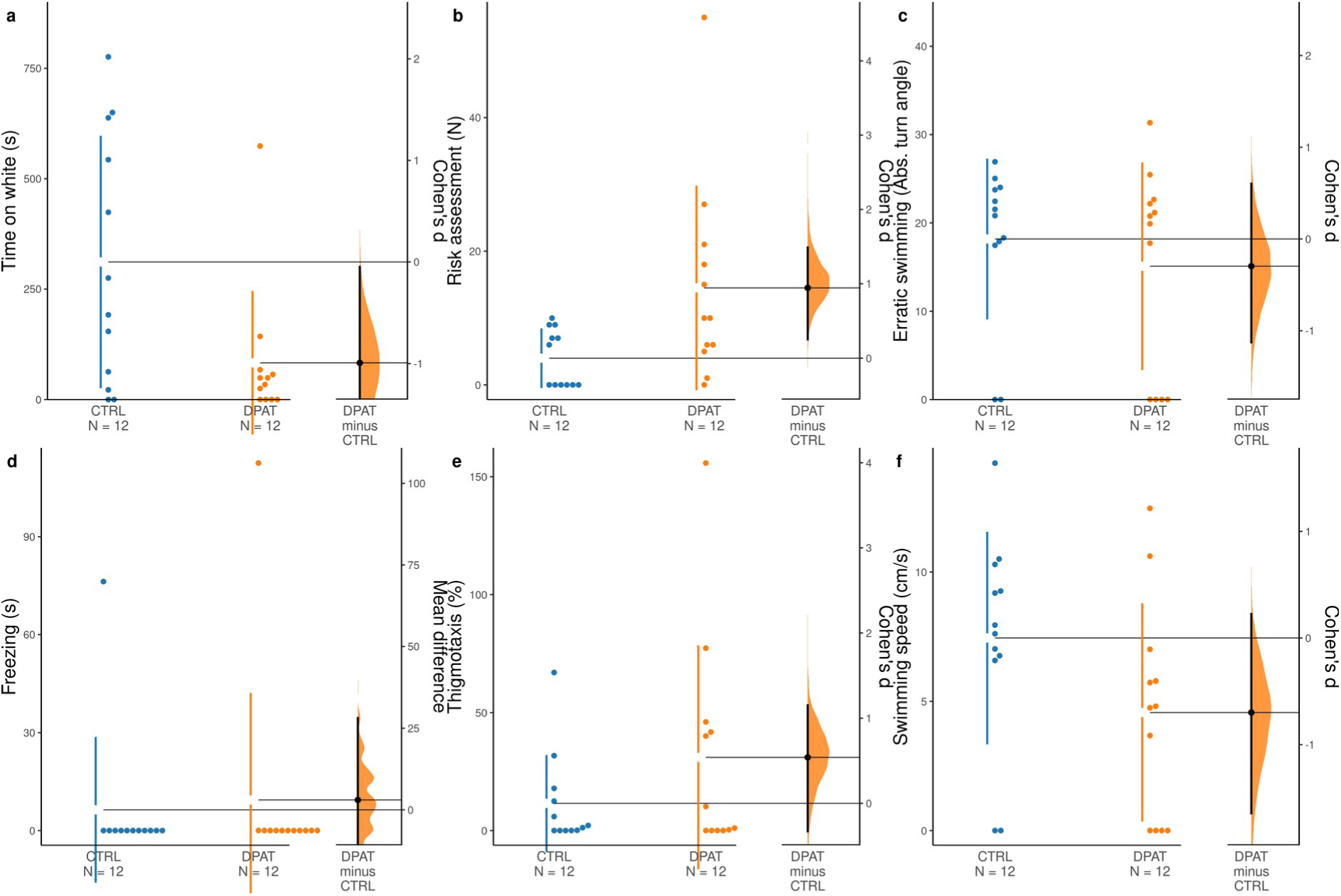
– 8-OH-DPAT increases anxiey-like behavior in the zebrafish light/dark test. (A) Scototaxis. (B) Risk assessment. (C) Erratic swimming. (D) Freezing. (E) Thigmotaxis. (F) Swimming speed. For a description of variables, refer to Table 1. Blue dots represent animals treated with vehicle (CTRL), and orange dots represent animals treated with 8-OH-DPAT (DPAT). The Cohen’s *d* between CTRL and DPAT is shown in Gardner-Altman estimation plot. Both groups are plotted on the left axes; the effect size (Cohen’s *d*) is plotted on floating axes on the right as a bootstrap sampling distribution. The mean difference is depicted as a dot; the 95% confidence interval is indicated by the ends of the vertical error bar. 5000 bootstrap samples were taken; the confidence interval is bias-corrected and accelerated.

### 3.3. Social preference test

During the SI phase, 8-OH-DPAT decreased the preference score (t_[df = 13.466]_ = 2.981, p = 0.0103, power = 0.81; Figure 3A), with no effects on social avoidance (t_[df=21.594]_ = 0.881, p = 0.388, power = 0.13; Figure 3B). No effects were observed on erratic swimming (t_[df = 21.392]_ = 0.548, p = 0.589, power = 0.08; Figure 3C), nor on swimming speed (t_[df = 19.036]_ = 1.1803, p = 0.252, power = 0.2; Figure 3D). For all variables, 95% confidence intervals for the effect sizes for sex X group interactions overlapped zero (Supplemental File S3); however, plots suggest no differences in preference score between males and females, and larger effects of 8-OH-DPAT in females vs. male zebrafish in this variable; likewise, no differences are suggested in social avoidance, but larger effects of 8-OH-DPAT in males than females are suggested. Plots also suggest that females show higher erratic swimming than males, and 8-OH-DPAT appears to reduce erratic swimming in males but not females in the SI phase.

**Figure 3.**
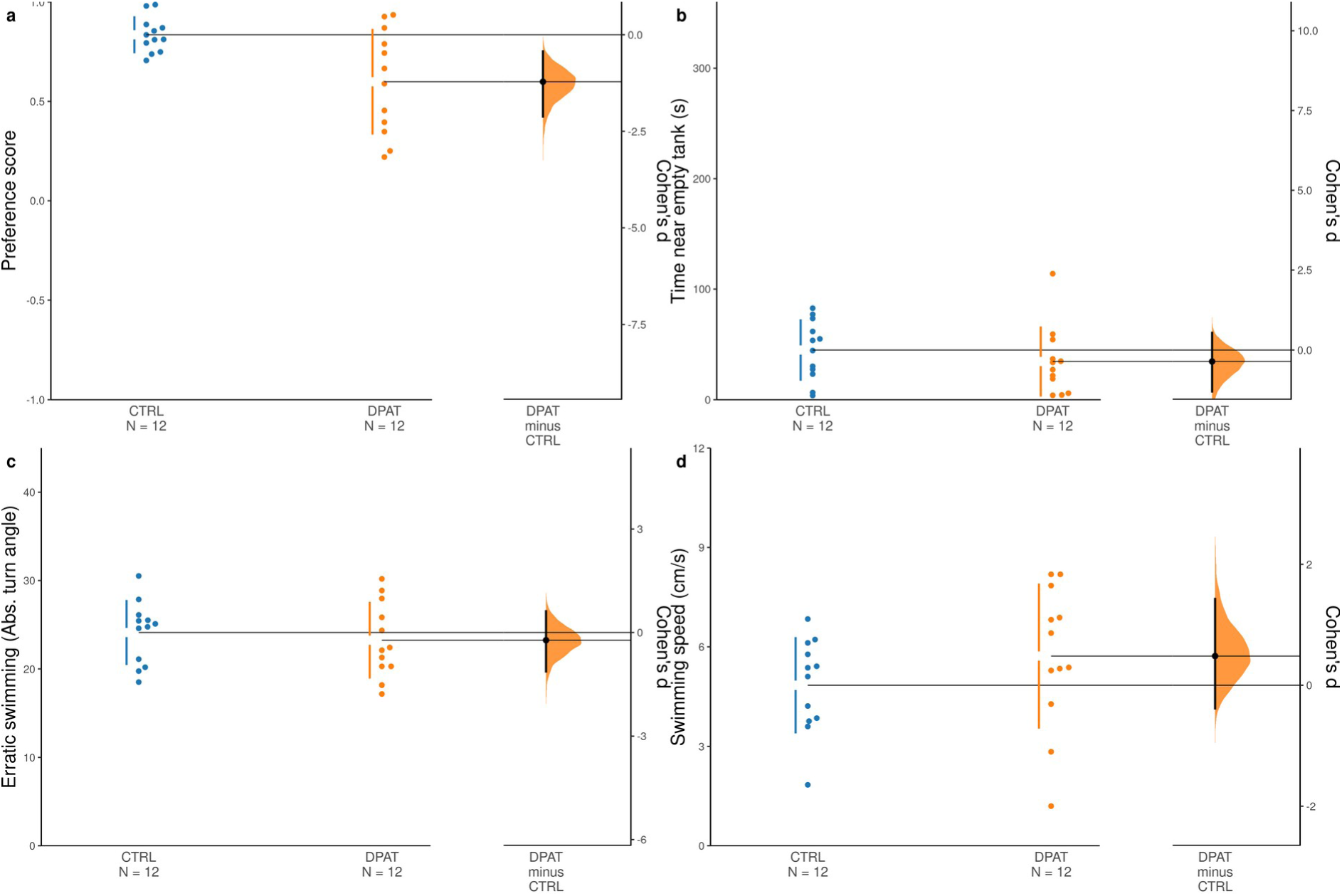
– 8-OH-DPAT decreases social approach in the social investigation test. (A) Preference score. (B) Social avoidance. (C) Erratic swimming. (D) Swimming speed. For a description of variables, refer to Table 1. Blue dots represent animals treated with vehicle (CTRL), and orange dots represent animals treated with 8-OH-DPAT (DPAT). The Cohen’s *d* between CTRL and DPAT is shown in Gardner-Altman estimation plot. Both groups are plotted on the left axes; the effect size (Cohen’s *d*) is plotted on floating axes on the right as a bootstrap sampling distribution. The mean difference is depicted as a dot; the 95% confidence interval is indicated by the ends of the vertical error bar. 5000 bootstrap samples were taken; the confidence interval is bias-corrected and accelerated.

During the SN phase 8-OH-DPAT also decreased the preference score (t_[df = 21.847]_ = 2.667, p = 0.014, power = 0.72; Figure 4A), but the time near the “new stranger” was also decreased (t_[df = 21.767]_ = 2.701, p = 0.013, power = 0.73; Figure 4B). No effect was observed on erratic swimming (t_[df = 21.785]_ = −0.7101, p = 0.485, power = 0.1; Figure 4C), nor on swimming speed (t_[df = 21.797]_ = −1.359, p = 0.188, power = 0.25; Figure 4D). For all variables, 95% confidence intervals for the effect sizes for sex X group interactions overlapped zero (Supplemental File S4); again, plots suggest no differences in preference score between males and females, and larger effects of 8-OH-DPAT in males vs. female zebrafish in this variable; likewise, no differences are suggested in social avoidance, but larger effects of 8-OH-DPAT in males than females are suggested. Plots also suggest that females show higher erratic swimming than males, and 8-OH-DPAT appears to reduce erratic swimming in females but not males in the SN phase.

**Figure 4.**
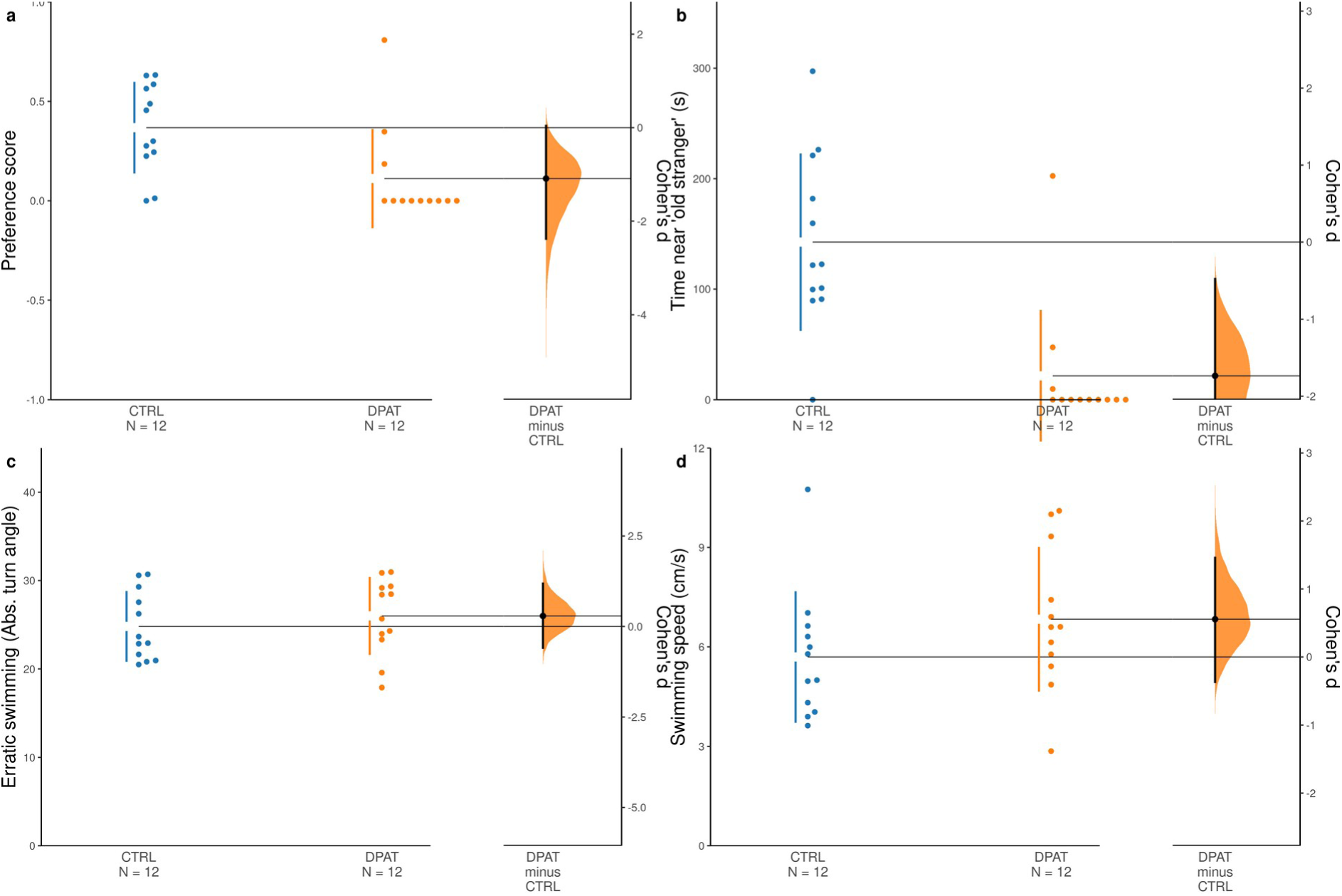
– 8-OH-DPAT decreases social preference in the social novelty test. (A) Preference score. (B) Social avoidance. (C) Erratic swimming. (D) Swimming speed. For a description of variables, refer to Table 1. Blue dots represent animals treated with vehicle (CTRL), and orange dots represent animals treated with 8-OH-DPAT (DPAT). The Cohen’s *d* between CTRL and DPAT is shown in Gardner-Altman estimation plot. Both groups are plotted on the left axes; the effect size (Cohen’s *d*) is plotted on floating axes on the right as a bootstrap sampling distribution. The mean difference is depicted as a dot; the 95% confidence interval is indicated by the ends of the vertical error bar. 5000 bootstrap samples were taken; the confidence interval is bias-corrected and accelerated.

### 3.4. Effects of 8-OH-DPAT after WAY 100,635 pre-treatment

Main effects of 8-OH-DPAT dose (F_[3, 72]_ = 111.55, p < 0.001, ω² = 0.68, power = 0.99) and WAY 100,635 dose (F_[1, 72]_ = 32.73, < 0.0001, ω² = 0.06, power = 0.46) were found for geotaxis. Importantly, an interaction effect was also found (F_[3, 72]_ = 16.71, p < 0.001, ω² = 0.1, power = 0.15; Figure 5). Post-hoc tests found that, when animals were not pre-treated with WAY 100,635, 8-OH-DPAT decreased geotaxis at all doses (p < 0.05, *d* between 1.42 and 5.25); however, in animals that were pre-treated with WAY 100,635, 8-OH-DPAT produced a significant reduction only at the highest dose (p < 0.001, *d* = 5.17).

**Figure 5.**
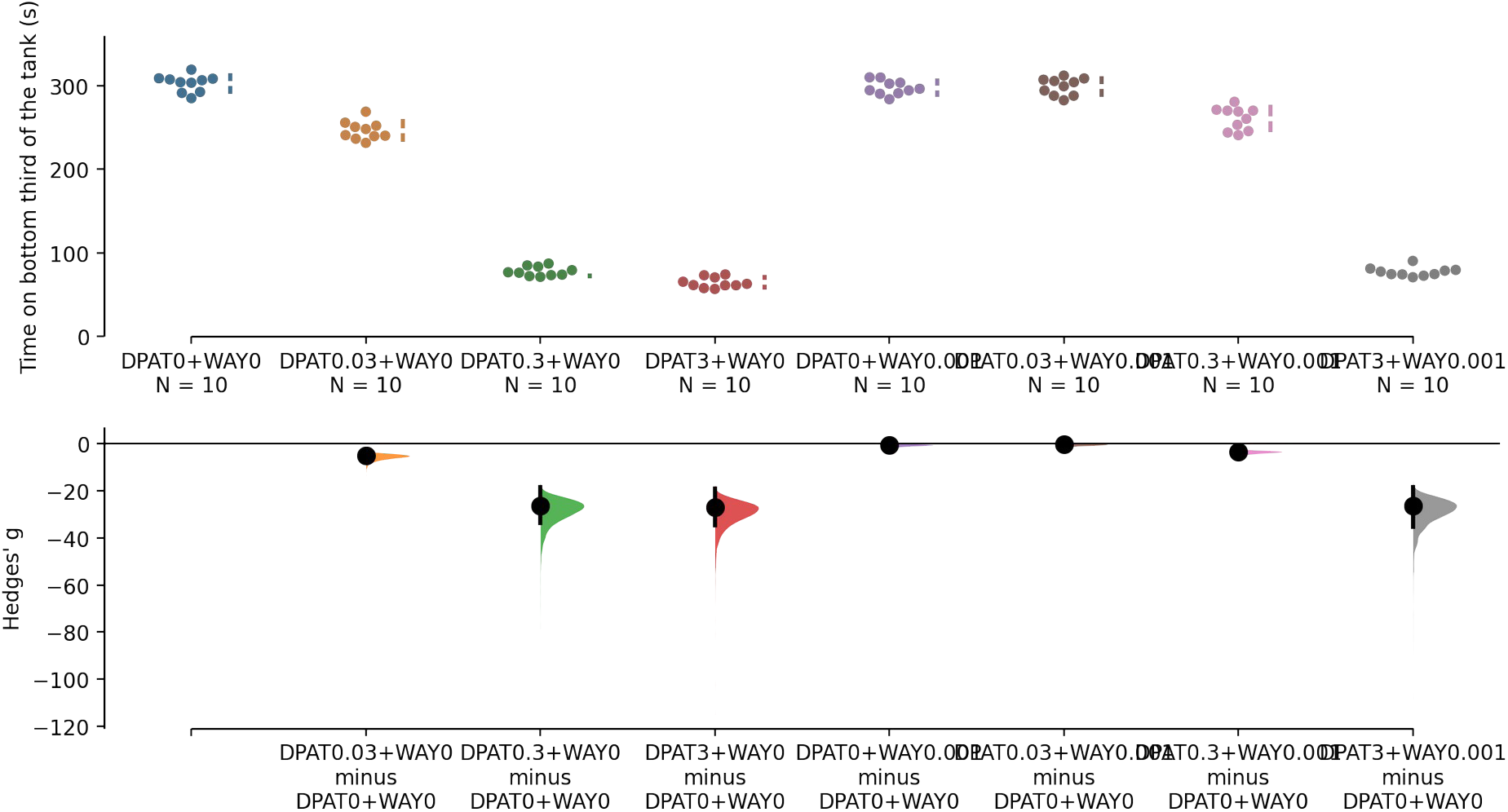
– The Hedges’ *g* for 7 comparisons against the shared control DPAT0+WAY0 are shown in a Cumming estimation plot. The raw data is plotted on the upper axes. On the lower axes, mean differences are plotted as bootstrap sampling distributions. Each mean difference is depicted as a dot. Each 95% confidence interval is indicated by the ends of the vertical error bars.

## 4. Discussion

In the present experiments, differences were observed in anxiety-like behavioral endpoints in the groups treated with the full agonist of the 5-HT1A receptor, 8-OH-DPAT, with opposite outcomes in the novel tank test (NTT) and phototaxis assay (light-dark test). While in the first 8-OH-DPAT decreased anxiety-like behavior, in the latter the drug increased it. When 8-OH-DPAT was injected in animals pre-treated with WAY 100,635, a rightward shift in the dose-response curve was found in the NTT. We also observed decreases in social preference in both the social interaction (SI) and social novelty (SN) tests.

### 4.1. Effects of 8-OH-DPAT on anxiety tests

One of the most striking findings of this study is the difference in the effects of 8-OH-DPAT between the phototaxis assay and the NTT. In previous studies with serotonergic drugs in these two tests, it was shown that increasing 5-HT levels with acute fluoxetine increases geotaxis (the main endpoint of NTT) and decreases scototaxis (the main endpoint of the phototaxis assay), while decreasing 5-HT levels with pCPA produces the opposite effects (Maximino et al., 2013). On the other hand, both buspirone (partial 5-HT1AR agonist) and WAY 100,635 (full 5-HT1AR antagonist) decreased geotaxis and scotototaxis (Maximino et al., 2013). SB 224,289, an inverse agonist of the 5-HT1B receptor, reduced geotaxis but not scototaxis, an effect similar to that observed in the present experiment. Unlike buspirone, which produces consistent reductions in responses to potential threat in both the phototaxis assay and the NTT (Abozaid & Gerlai, 2022; Araujo et al., 2012; Bencan et al., 2009; Connors et al., 2014; Facchin et al., 2015; Gebauer et al., 2011; Hawkey et al., 2021; Lau et al., 2011; Maaswinkel et al., 2012, 2013; Maximino et al., 2011, 2013, 2014; Steenbergen et al., 2011), 8-OH-DPAT seemed to produce opposite effects in the NTT and phototaxis assay. Despite being 5-HT1A receptor agonists, there are important differences between buspirone and 8-OH-DPAT. In a context of large receptor reserves in presynatic neurons, as is the case with the rodent raphe, buspirone acts as a full agonist at the 5-HT1A autoreceptor, while it acts as an antagonist at the postsynaptic 5-HT1AR, reducing serotonin avaiability at other postynapic sites while simultaneously blocking postsynaptic 5-HT1ARs (Newman-Tancredi et al., 2019). Meanwhile, 8-OH-DPAT is a serotonergic drug that acts as a full agonist at both sites, reducing serotonin availability for other types of receptor in the postsynaptic sites while simultaneously activating postsynaptic 5-HT1ARs. It is therefore possible to assume that this is also the reason why the results differed between the tests.

To confirm the presence of a receptor reserve related to the effects of 8-OH-DPAT, animals were pre-treated with the antagonist WAY 100,635 and treated with increasing doses of 8-OH-DPAT before being subjected to the NTT. A rightward shift in the dose-response curve was observed, with higher doses needed to produce an anxiolytic-like effect in the NTT when receptors were blocked. When rats are treated with 8-OH-DPAT and the irreversible 5-HT1A antagonist EEDQ, a rightward shift is observed in the production of 5-HTP, the serotonin precursor, in the cortex and hippocampus (Meller et al., 1990). Likewise, Cox et al. (1993) showed that EEDQ induced a rightward shift in the dose-response curve for the inhibition of dorsal raphe nucleus cell firing by 8-OH-DPAT. Although these studies were the first to show evidence of a receptor reserve for neurophysiological endpoints of serotonergic function, other studies found mixed evidence for systemic endpoints. For example, Meller and Bohmaker (1994) found that EEDQ induces a rightward shift in the attenuation of plasmatic release of corticosterone and ACTH in rats. However, no evidence for receptor reserve was found for 5-HT1A receptor-mediated hypothermia in mice (Meller et al., 1992). Overall, these results are consistent with the hypothesis that, at least for behavior in the NTT, a receptor reserve is present.

A related question is why 8-OH-DPAT reduced behavioral responses to potential threat in the NTT and increased it in the LDT. One possibility is that NTT is more sensitive to pharmacological treatment than LDT. However, in a meta-analysis of the effects of drugs on the two tests, Kysil et al. (2017) found no differences in the sensitivity of both assays. A second possibility is that the effect observed on the NTT is not related to a decrease in responsiveness to potential threat, but represents either serotonin toxicity (Stewart et al., 2013) or a psychedelic-like effect (Kyzar & Kalueff, 2016). While 8-OH-DPAT increased top-dwelling in the NTT, which could be interpreted as “surfacing” (an aberrant behavioral pattern of slow swimming close to the water surface that is typically observed under the effects of serotonergic hallucinogens; Kalueff et al., 2013; Kyzar & Kalueff, 2016), animals treated with the drug still spent most of the time in the bottom of the tank, differently from zebrafish treated with psychedelics. Moreover, psychedelics typically produce an hypolocomotion phenotype in zebrafish, and changes in the swimming speed were not observed in any test in the present experiments. A third possibility is that 8-OH-DPAT is acting on either 5-HT1A and 5-HT7 receptors, producing different behavioral profiles depending on the receptor. While the affinity of this drug for zebrafish receptors has not been determined, it could be argued that the opponent effects of 8-OH-DPAT on the NTT and the phototaxis assay are due to effects on these receptors. The scant amount of behavioral studies with 8-OH-DPAT investigating these receptors make it difficult to judge that interpretation. In rodents, 8-OH-DPAT induces hypothermia by acting both on the 5-HT1A and the 5-HT7 receptors (Hedlund et al., 2004). However, 5-HT7R knockout mice showed no differences in baseline anxiety (Guscott et al., 2005), suggesting that this receptor is involved in other behavioral functions, at least in mammals. To fully confirm this hypothesis, more experiments are needed.

An alternative explanation is that the two tests model different defensive processes, with opposite modulation of the serotonergic system on both. Blaser et al. (2012) analyzed stimulus control in the two tests, and suggested that geotaxis is mainly controlled by escape from the top (closer, therefore, to fear), while scototaxis is mainly controlled by approach-avoidance conflict (closer, therefore, to anxiety). Thus, the effects of 8-OH-DPAT in the NTT and phototaxis assay could be interpreted as decreasing fear and increasing anxiety, respectively. In support of this hypothesis, microinjection of 8-OH-DPAT in forebrain regions such as the hippocampus (Cheeta et al., 2000; File et al., 1996), septum (Cheeta et al., 2000; Martins De Almeida et al., 1998), and amygdala (Nunes-de-Souza et al., 2000) has been found to increase anxiety-like behavior in rodents. Conversely, microinjection of 8-OH-DPAT in the periaqueductal gray (Zangrossi Jr. et al., 2001; Zanoveli et al., 2003, 2010) and dorsomedial hypothalamus (Nascimento et al., 2014), but not amygdala (Zangrossi et al., 1999), impair escape in the elevated T-maze, an effect consistent with decreased fear-like states. In zebrafish, treatment with the 5-HT1A antagonist WAY 100,635 increases responsiveness to fear-eliciting olfactory stimulus (alarm substance), implicating this receptor in reducing responses to actual threats (Nathan et al., 2015). Taken together, these results suggest that the differential effects of 8-OH-DPAT on the NTT and phototaxis assay mediate different behavioral processes via 5-HT1A heteroreceptors. Experiments using 8-OH-DPAT to assess its effects on alarm substance-elicited behavioral responses could test this hypothesis. Considering the participation of the septal homologue Vv and the hypothalamic Hv in scototaxis (Lau et al., 2011), and considering that both 5-HT1AR isoforms are found in the latter while only *htr1aa* is found in Vv (Norton et al., 2008), is is likely that these nuclei are involved in the effects of 8-OH-DPAT.

A last possibility that cannot be discarded is that 8-OH-DPAT acts on receptors involved with locomotion, either at the hindbrain or spinal cord levels; indeed, *htr1aa* has been found to be expressed in hindbrain motor circuits (Norton et al., 2008). In zebrafish larvae, treatment with methysergide, a non-selective agonist of 5-HT1 receptors and antagonist at 5-HT2 receptors, decreases motor output by prolonging periods of inactivity (Brustein et al., 2003). Thus – especially in the NTT, in which the main endpoint involves the ability to swim upwards in the water column –, this motor-inhibitory effect of 5-HT1A receptor activation could be responsible for the observed effects. While the lack of effects on swimming speed on all tests weakens this hypothesis, post-hoc power analysis showed that this variable had lower power than desired across assays, and therefore interpretations on these effects should be taken with a grain of salt.

In the present experiments, no effects were observed in measures such as erratic swimming, freezing, and thigmotaxis, which are common secondary endpoints in the NTT and phototaxis assay. Previously, it has been shown that WAY 100,635 and buspirone decreased freezing, swimming, and thigmotaxis in the phototaxis assay at higher doses, but had no effect in these endpoints in the NTT (Maximino et al., 2013). Nowicki et al. (2014), on the other hand, found that m-PPF, another 5-HT1AR antagonist, increased erratic swimming at low doses in the NTT. Post-hoc power analyses also showed that these variables had lower power than desired, suggesting that the lack of effects in the present experiments in these variables are likely to be false negatives.

### 4.2. Effects of 8-OH-DPAT on social approach

Different phases of the social preference tests used in the present experiment emphasize the role of social interaction vs. social novelty. Our results with 8-OH-DPAT showed a decrease in SI and SN, suggesting a broader effect on social approach. Barba-Escobedo and Gould (2012) showed that buspirone increases social approach in the SI test (the opposite of what was observed here with 8-OH-DPAT), with no effect on the SN test. Whether this difference indicates that, as in the case with anxiety-like behavior, the effects of 8-OH-DPAT are likely to be mediated by heteroreceptors is still opened. These results could be interpreted either as an anxiogenic-like effect on social preference, or a specific effect on sociality. These hypotheses are difficult to untangle, since, while no effects on erratic swimming was observed in neither the SI or SN tests, they were also absent in the NTT or phototaxis assay. While social approach is sensitive to acute stress in adult zebrafish (Cook et al., 2023), the fact that, differently from lab rodents, zebrafish do not avoid novel conspecifics, and actively spend time investigating them when possible (Barba-Escobedo & Gould, 2012; do Nascimento & Maximino, 2023; Ogi et al., 2021), makes it difficult to understand how threat factors in sociability in this species. The activation of 5-HT2 receptors has been shown to increase sociability in zebrafish (de Moura et al., 2023; Ponzoni et al., 2016), and single-nucleotide polymorphisms of the *htr1aa* gene (which code for one isoform of the 5-HT1AR) mediate populational variability in social exploration and sociability in this species (Gonçalves et al., 2022). These results further underscore the role of serotonin in sociability in zebrafish, with opponent modulation by 5-HT1AR and 5-HT2CR agonists.

As was the case with the LDT and NTT, no effects were observed in erratic swimming. This can suggest that the effects of 8-OH-DPAT are restricted to sociality, and not an increase in anxiety-like behavior. However, as was the case with the LDT and NTT, post-hoc power analysis showed lower power than desired, and therefore it is difficult to understand whether or not the lack of effect in erratic swimming in the SI and SN tests are false negatives.

### 4.3. Exploration of potential sex differences

While the current study was not designed to detect sex differences and interactions between sex and drug effects, some suggestion of such effects were observed. In the NTT, a small effect sizes of sex were observed in both geotaxis and top-dwelling, with females displaying less geotaxis and more top-dwelling. While exploratory, these results contradict what is usually reported in the literature (Genario et al., 2020). Moreover, moderate effect sizes on the interaction are also suggested, with females showing a possibly higher effect of 8-OH-DPAT than males. In the phototaxis assay, effects appear to be opposite, with higher anxiety-like behavior in females (small effect size) and a higher effect of 8-OH-DPAT in males (moderate effect size). Thus, important sex effects could also be responsible for the opposite directions of 8-OH-DPAT effects in the NTT and phototaxis assay; experiments explicitly designed to test this hypothesis are needed. In the SI phase of the social preference test, almost no difference is obserable between males and females, but larger effects of 8-OH-DPAT in females vs. male zebrafish are suggested in social approach, as are larger effects of 8-OH-DPAT in males than females in social avoidance. Similar effects were observed in the SN phase, but larger effects of 8-OH-DPAT in males than females are suggested. These results are consistent with previous work which demonstrated that both males and females show similar social preferences (Snekser & Diestler, 2023). We emphasize, again, that these analyses are exploratory, and sample sizes were not planned to detect main or interaction effects of sex. Thus, further experiments are necessary to confirm these hypotheses of sex effects.

### 4.4. Limitations

One important limitation of the present study is that, although sample sizes were higher than those calculated as the minimum to produce an effect size at least as large as that of buspirone, post-hoc power analyses found that, for important secondary endpoints, power was lower than desired. Due to IRB restrictions on increasing sample sizes after the study was conducted, it was unfeasible to re-run the experiments to confirm whether 8-OH-DPAT did not produce an effect on erratic swimming and freezing, or if the apparent lack of effect is a false negative due to low power. Further experiments are necessary to untangle this hypothesis.

Another important limitation relates to how to accurately interpret the results of the social preference tests in zebrafish. In rats, decreases in social interaction is usually interpreted as indicating an anxiogenic-like effect (File & Seth, 2003). In zebrafish, some studies suggest that acute stress decreases social approach (Barcellos et al., 2020; Cook et al., 2023); nonetheless, while buspirone increases social approach (Barba-Escobedo & Gould, 2012), non-serotonergic anxiolytics such as diazepam appear to decrease it in zebrafish (Giacomini et al., 2016). Future work should also disentangle the participation of threat (and therefore of anxiety-like states) and sociability in this test, allowing for a better comprehension of the roles of the 5-HT1AR in it.

A third limitation of the present study is that it was not explicitly design to take into account possible sex differences both in the behavioral parameters and in the drug effects. There is reason to believe that there are differences in the serotonergic system of male and female zebrafish, as a lower 5-HIAA:5-HT ratio was observed in the forebrain of females (Dahlbom et al., 2012). Our exploratory analysis suggested that males and females respond differently to 8-OH-DPAT, and therefore studies explicitly designed to address this hypothesis are needed.

Finally, our study did not include an explicit dose-response analysis, as only one dose of 8-OH-DPAT was used. Thus, it is possible that the dose used was not optimal for the phototaxis assay and the social preference tests. While this could also explain the lack of effects of 8-OH-DPAT on secondary endpoints, such as freezing and erratic swimming, it is also important to point that dose-response curves under concomitant treatment with receptor antagonists are commonly used to assess the presence of a receptor reserve (Cox et al., 1993; Furchgott & Bursztyn, 1967; Meller et al., 1990). While dose-response curves in behavioral pharmacology are usually different from the typical sigmoidal curve found elsewhere (e.g., lower doses of WAY 100,635 produce larger effects in the NTT and LDT than higher doses; Maximino et al., 2013), it can be useful to perform such experiments in the future to better understand whether a receptor reserve is responsible for the different effects of 8-OH-DPAT and other 5-HT1AR binding drugs in these behavioral tests.

### 4.5. Translational implications

The present work shed light on a serotonergic mechanism involved in the opponent regulation of responses to proximal threat in zebrafish. While substantial differences exist between zebrafish and humans in terms of their serotonergic system (Herculano & Maximino, 2014), the fact that 8-OH-DPAT appears to produce similar behavioral effects in both zebrafish and rodents increase the possibility that the function of 5-HT1ARs is conserved. Indeed, it has been previously proposed that using non-mammalian organisms in behavioral neuroscience can facilitate a deeper understanding of brain function and dysfunction at different levels (Gerlai, 2020; Stewart et al., 2014). If similar drug effects are found in zebrafish, rodents, and humans, it is likely that those effects represent conserved physiology instead of evolutionary homoplasies (Gerlai, 2020). In the context of molecular heterogeneity and physiological conservation of the serotonergic system, the present results point to the importance of interpreting data from behavioral pharmacology in a comparative framework.

## 5. Conclusions

In the present experiments, the full agonist of the 5-HT1A receptor, 8-OH-DPAT, decreased anxiety-like behavior in the novel tank test (NTT), but increased it in the light-dark test (LDT), both considered outcomes of anxiety-like behavior for this species. The same treatment decreased social approach in both the social investigation test and the social novelty test. In comparison with previous data on the effects of the partial agonist buspirone and the full antagonist WAY 100,635, and based on the evidence for a receptor reserve for the anxiolytic-like effect of 8-OH-DPAT in the NTT, we suggest that 8-OH-DPAT increases anxiety-like behavior mainly by activating heteroreceptors.

## Acknowledgments

This work was supported by a Conselho Nacional de Desenvolvimento Científico e Tecnológico (CNPq/Brazil) productivity grant to CM (#302998/2019-5). LVXBS an LNO were supported by Fundação Coordenação de Aperfeiçoamento de Pessoal de Nível Superior (CAPES/Brazil) scholarships. A GPT-3.5-based tool, SciSummary, was used to help create the lay summary and the significance statement of this article, but no other AI-based tools were used elsewhere. The graphical abstract for this manuscript was created with BioRender.com

## Conflict of interest statement

The authors declare that they have no conflicts of interest.

## Authors’ contributions

**Loanne Valéria Xavier Bruce de Souza:** Conceptualization, Methodology, Formal Analysis, Investigation, Writing – Original draft, Writing – Review & editing, Visualization; **Larissa Nunes de Oliveira:** Investigation, Writing – Original draft, Writing – Review & editing; **Bruna Patrícia Dutra Costa:** Investigation, Writing – Original draft, Writing – Review & editing; **Monica Lima-Maximino:** Conceptualization, Methodology, Writing – Original draft, Writing – Review & editing; **Hellen Vivianni Veloso Corrêa:** Conceptualization, Methodology, Writing – Original draft, Writing – Review & editing; **Caio Maximino:** Conceptualization, Methodology, Validation, Formal Analysis, Resources, Data Curation, Writing – Original draft, Writing – Review & editing, Supervision, Project administration, Funding acquisition.

## Data accessibility

The data that supports the findings of this study are openly available in GitHub/Figshare at https://doi.org/10.6084/m9.figshare.26272720

## Notes

https://github.com/lanec-unifesspa/5HT/tree/main/5HT1A

### Competing Interest Statement

The authors have declared no competing interest.

### Summary of Updates

New results included, from an experiment on receptor reserve

